# Glycosylation-associated dysregulation of pyocyanin production in *Pseudomonas aeruginosa*: Implications for quorum sensing regulation

**DOI:** 10.1101/319418

**Authors:** Anna K. McClinton, Caleb L. Hamilton, Donna L. Cioffi, Eugene A. Cioffi

## Abstract

*Pseudomonas aeruginosa* (*P. aeruginosa*) is an important opportunistic pathogen associated with high mortality in pneumonia, sepsis, and cystic fibrosis. Lending to its ability to cause severe disease and death is its arsenal of virulence factors and host evasion tactics. In addition to various other regulatory systems, many of *P. aeruginosa*’s virulence factors are regulated by a population density dependent regulatory network known as quorum sensing (QS). Many regulatory systems are impacted by post-translational modifications of proteins. An underexplored physiological aspect of *P. aeruginosa* is its ability to glycosylate proteins and the subsequent impact of glycosylation on *P. aeruginosa* physiology and behavior. The goal of this study was to determine whether *P. aeruginosa* QS is regulated by glycosylation. Here we demonstrate that disruption of glycosylation dysregulates QS phenotypes, notably pyocyanin production, in *P. aeruginosa* PAO1. In this study, it was initially observed that deletion of the *P. aeruginosa* neuraminidase, PaNA, caused an increased production of pyocyanin in LB-Lennox broth compared to wildtype bacteria at identical population densities. To confirm that the increased pyocyanin production was due to QS, we performed induction experiments using 10% cell-free media harvested from overnight cultures. To determine whether the QS phenotype observed is specific to pseudaminic acid, the target of PaNA, or if it is a reflection of global changes in glycosylation, we measured QS in a library of mutant bacteria generated in an MPAO1 background containing transposon insertions in various glycosyl-associated enzymes. The pattern of dysregulated QS held true in these mutant strains as well. Overall these data indicate that in *P. aeruginosa*, glycosylation is an important determinant of QS.

## Introduction

*P. aeruginosa* is a gram-negative bacteria, found ubiquitously in the soil and water[1, 2]. It is well suited to exploiting a broad range of environments including human hosts. In humans, *P. aeruginosa* is associated with acute conditions such as burn wound infections, eye infections, and ventilator-associated pneumonia and sepsis, as well as chronic infections of the cystic fibrosis lung. Despite its being an opportunistic pathogen, *P. aeruginosa* can cause severe and fatal infections. *P. aeruginosa* infection is multifactorial as the bacteria possess a plethora of virulence factors. An aspect of the multifactorial virulence of *P. aeruginosa* is that the regulation of virulence factors is highly complex and mutli-layered[2]. One of the layers, quorum sensing (QS), is accountable for the control of several hundred genes, many of which relate to virulence[3]. QS is a phenomenon of bacterial communication and coordination which is traditionally defined as dependent on population density, whereby when a small molecule, generated at a steady rate corresponding with population density, is accumulated at sufficient quantities, a complex regulatory cascade is triggered in the population controlling the expression and repression of several hundred genes[4, 5]. This complex autoinduction system has at least three separate, yet interdependent arms: the Las system, the Rhl system and the PQS system[5-7]. Each of these systems is complex and capable of cross-talk[7-9]. QS is one example of *P. aeruginosa* responding to its external environment and the regulation of this phenomenon is dynamic and complex. A number of upstream signal transduction systems have been implicated in the regulation of quorum sensing[10-13]. One post-translational modification that is unexplored in QS regulation is glycosylation.

Signal transduction occurs across the bacterial membrane, as does glycosylation[13, 14]. Glycosylation is the covalent attachment of a glycan, or carbohydrate chain, to a substrate such as a protein or a lipid[15, 16]. While protein glycosylation was previously viewed as a eukaryotic process, it is now recognized as a process that occurs across all domains of life[16]. In *P. aeruginosa,* only a few proteins have been identified as glycosylated and these modifications do not necessarily occur in all strains[17]. Known examples of glycosylation in *P.* aeruginosa include the flagella, pili, and LecB lectin[14, 17-28]. While much remains unknown concerning protein glycosylation in *P. aeruginosa*, a great deal has been elucidated concerning lipid glycosylation. Interestingly, the biosynthetic pathways of LPS correlate or even overlap with the few known pathways of protein glycosylation[29-31]. The majority of the enzymes involved in LPS biosynthesis have been characterized, however, uncharacterized enzymes predicted to be glycosyltransferases remain [29, 30]. In this study we observed an altered QS phenotype in a small library of bacteria carrying transposon insertions in various, “probable” glycosyl-associated enzymes[32-34], providing a link between glycosylation and QS. These glycosyl-associated enzymes encompass glycosyl hydrolases which would be responsible for removing carbohydrates from a glycan chain by hydrolysis as well as glycosyl-transferases which add to individual carbohydrates to glycan chains.

An interesting carbohydrate found in eukaryotes and bacteria is sialic acid. Sialic acids are any of the 9 carbon, α-keto sugars derived from neuraminic acid[35]. Sialic acids often occur as the terminal sugar of the carbohydrate branches of glycans[36]. A specialized sialic acid-like sugar known as pseudaminic acid is found in some bacteria and has been identified as a component of the glycan found on pilin of some strains of *P. aeruginosa.* Pseudaminic acid has been found decorating the flagella of several other gram negative bacteria, as well[37]. Additionally, pseudaminic acid has been found as component of LPS in a variety of gram negative bacteria and in a few strains of *P. aeruginosa*[37-43]. Sialic acids are cleaved from the underlying glycan by a class of enzymes referred to as sialidases or neuraminidases[44]. *P. aeruginosa* PAO1 possesses an enzyme initially identified as a neuraminidase, but following X-ray crystallography and *in silico* docking experiments, the enzyme was determined to be a pseudaminidase[45, 46]. This enzyme, PaNA, is encoded at locus PA2794 and was initially of interest as a virulence factor. However, its role in the biology of the bacteria remains uncharacterized[36, 44].

A strain of *P. aeruginosa* PAO1 from which the neuraminidase gene PA2794 has been deleted[44], PAO1Δ2794, exhibited a pronounced over-expression of pyocyanin, which is associated with the PQS arm of QS, compared to the wildtype strain. We therefore hypothesized that the deletion of the neuraminidase resulted in an alteration of the glycosylation of one or more proteins which led to this anomalous phenotype. Indeed, lectin blots revealed a differential pattern of glycosylation between the wildtype strain and the mutant Δ2794. We confirmed our initial observations by using a different PaNA mutant, PW5679, generated in MPAO1 by transposon insertion. To determine whether the QS phenotype observed is specific to pseudaminic acid, the target of PaNA, or if it is a reflection of global changes in glycosylation, we measured QS in a library of mutant bacteria generated in an MPAO1 background containing transposon insertions in various glycosyl-associated enzymes. The pattern of dysregulated QS held true in these mutant strains as well. We show that disruption of PaNA, as well as other glycosyl-associated enzymes, results in a QS phenotype—namely the over-production of pyocyanin—which is decoupled from population density, overall suggesting that bacterial glycosylation is a critical determinant of QS.

## Materials & Methods

### Bacterial Strains and Growth Conditions

Table I presents the strains utilized in this study. Strains were routinely grown in LB-Lennox broth (Sigma-Aldrich; Darmstadt, Germany) and/or agar plates (BD Difco; Franklin Lakes, NJ). Freezer stocks were maintained in nutrient broth (BD Difco) with 12.5% glycerol (Sigma) at −80°C. Overnight cultures of the transposon mutants were grown in the presence of tetracycline.

**Table 1.**

| Strain Name | Description | Source |
| --- | --- | --- |
| PAO1 | Wildtype | (Soong et al., 2006) |
| PAO1Δ2794 | Allelic deletion of PA2794<br>Constructed in PAO1 background<br>Disruption of Pseudaminidase gene | (Soong et al., 2006) |
| PW2532 | PA0842-D03::ISlacZ/hah<br>Constructed in MPAO1 background<br>Disruption of a probable glycosyltransferase | (Held, Ramage, Jacobs, Gallagher, & Manoil, 2012; Jacobs et al., 2003) |
| PW2834 | PA1014-G02::ISlacZ/hah<br>Constructed in MPAO1 background<br>Disruption of a probable glycosyltransferase | (Held et al., 2012; Jacobs et al., 2003) |
| **PW3519** | PA1385-F08::ISlacZ/hah<br>Constructed in MPAO1<br>background<br>Disruption of a probable<br>glycosyltransferase | (Held et al., 2012; Jacobs et al., 2003) |
| **PW3525** | PA1389-E07::ISlacZ/hah<br>Constructed in MPAO1<br>background<br>Disruption of a probable<br>glycosyltransferase | (Held et al., 2012; Jacobs et al., 2003) |
| **PW3528** | PA1390-H10::ISphoA/hah<br>Constructed in MPAO1<br>background<br>Disruption of a probable<br>glycosyltransferase | (Held et al., 2012; Jacobs et al., 2003) |
| **PW3531** | PA1391-B01::ISlacZ/hah<br>Constructed in MPAO1<br>background<br>Disruption of a probable<br>glycosyltransferase | (Held et al., 2012; Jacobs et al., 2003) |
| **PW4697** | PA2160-G04::ISlacZ/hah<br>Constructed in MPAO1<br>background<br>Disruption of a probable<br>glycosyl hydrolase | (Held et al., 2012; Jacobs et al., 2003) |
| **PW4699** | PA2162-G09::ISlacZ/hah<br>Constructed in MPAO1<br>background<br>Disruption of a probable<br>glycosyl hydrolase | (Held et al., 2012; Jacobs et al., 2003) |
| **PW4702** | PS2164-C10::ISlacZ/hah<br>Constructed in MPAO1<br>background<br>Disruption of a probable<br>glycosyl hydrolase | (Held et al., 2012; Jacobs et al., 2003) |
| **PW4801** | PA2233-H07::ISlacZ/hah<br>Constructed in MPAO1<br>background<br>Disruption of a probable<br>glycosyltransferase (PslC) | (Held et al., 2012; Jacobs et al., 2003) |
| **PW5679** | PA2794-G06::ISlacZ/hah<br>Constructed in MPAO1<br>background<br>Disruption of Pseudaminidase<br>gene | (Held et al., 2012; Jacobs et al., 2003) |
| **PW6130** | pelF-F08::ISphoA/hah<br>Constructed in MPAO1<br>background<br>Disruption of a PA3059, PelF, | (Held et al., 2012; Jacobs et al., 2003) |
|  | cytosolic glycosyltransferase |  |
| **PW9106** | PA4819-A03::ISphoA/hah<br>Constructed in MPAO1<br>background<br>Disruption of a probable<br>glycosyltransferase | (Held et al., 2012; Jacobs et al., 2003) |
| **PW7022** | ArnC-H11::ISlacZ/hah<br>Constructed in MPAO1<br>background<br>Disruption of PA3553, ArnC,<br>glycosyltransferase | (Held et al., 2012; Jacobs et al., 2003) |
| **PW3601** | LasI-F07::ISlacZ/hah<br>Constructed in MPAO1<br>background<br>Disruption of PA1432, LasI | (Held et al., 2012; Jacobs et al., 2003) |
| **PW3598** | LasR-C01::ISlacZ/hah<br>Constructed in MPAO1<br>background<br>Disruption of PA1430, LasR | (Held et al., 2012; Jacobs et al., 2003) |
| **PW6880** | RhlI-D03::ISphoA/hah<br>Constructed in MPAO1<br>background<br>Disruption of PA3476, RhlI | (Held et al., 2012; Jacobs et al., 2003) |
| **PW6882** | RhlR-B10::ISlacZ/hah<br>Constructed in MPAO1<br>background<br>Disruption of PA347, RhlR | (Held et al., 2012; Jacobs et al., 2003) |
| **PW2812** | MvfR-G11::ISlacZ/hah<br>Constructed in MPAO1<br>background<br>Disruption of PA1003, mvfR | (Held et al., 2012; Jacobs et al., 2003) |
| **MPAO1** | Wildtype (background for<br>transposon mutants) | (Held et al., 2012; Jacobs et al., 2003) |

### Growth Curves

Growth curves were conducted by inoculating 100 ml of sterile LB broth with 1 ml of overnight cultures. Cultures were grown at 37°C with shaking at 260 rpm. Optical Density at 600nm (OD600) was measured over the prescribed time course. Culture supernatant was removed and stored at −20°C for pyocyanin measurements. Pellets for use in lectin blots were stored at −80°C.

### Induction Bioassay

Bacteria were grown overnight (14-18 hours) in 100 ml of sterile LB broth with 1 ml of overnight cultures. Cultures were grown at 37°C with shaking at 260 rpm. Supernatant was harvested by two centrifugations of 45 minutes at 4000 rpm. The first centrifugation pelleted the bacteria, and the supernatant was transferred to fresh 50 ml conical tubes for the second centrifugation to clarify the supernatant further. The clarified supernatant was filter sterilized using 0.2 μm PES filters (Thermo Fisher Scientific; Waltham, MA). The filtrates were then aliquoted and stored at −20°C. Filtrates were thawed at 4°C overnight for use in the induction broth. Induction broth was prepared by adding 10% cell-free supernatant to naïve LB broth to a total volume of 100 ml. Prior to inoculation with 1 ml of overnight culture, 1 ml aliquots were taken from each flask to be utilized as a blank for pyocyanin measurements.

### Pyocyanin Measurement

Pyocyanin was measured over the course of the growth curves by collecting 4 ml aliquots and centrifuging at 4000 rpm for 20 minutes. The clarified supernatant was stored at −20°C. Pyocyanin was measured in clear 96-well Costar plates (Corning, Corning, NY) at an absorbance of 655 nm.

### Rhamnolipid Assessment

Swarming behavior was assayed as an indicator of rhamnolipid production. Swarm plates were poured at an agar concentration of 0.6% and allowed to equilibrate overnight at room temperature. Swarm plates were inoculated using sterile wooden picks and incubated inverted at 37°C in a humidified chamber overnight. Images of the plates were acquired and colonies outlined using ImageJ. The measurements were normalized to the area of the plate.

### Measurement of LasB Production

Skim-milk plates were utilized to assess LasB production. Skim-milk plates were poured by adding 20% sterile skim-milk to sterile, molten nutrient agar[47]. Plates were inoculated using sterile wooden picks and incubated, inverted at 37°C overnight. Images were acquired, and ImageJ was used to measure the area of the colonies and the area of the cleared zone surrounding the colonies. The measurements were normalized to the area of the plate and reported as the ratio of the zone to the colony.

### Lectin blots

Lectin blots were conducted on lysates from bacterial pellets harvested and stored at −80°C. Lysates were prepared as previously described[14] with modification, namely Triton-X (Sigma) was used in place of lysozyme. Protein was measured in the lysates using the BCA kit from Pierce (Thermo Fisher Scientific). Lysates were stored at −20°C. Lectin blots were performed by transferring the proteins to nitrocellulose membrane following SDS-PAGE in a 4-12% NuPAGE gel (Thermo Fisher Scientific). Ponceau staining of the membrane was performed to confirm equal protein loading. Membranes were blocked in 1X Carbofree buffer (Vector Laboratories; Burlingame, CA). Lectin reactivity was assessed with biotinylated Lotus lectin (Vector) was along with Streptavidin, DyLight 488 (Thermo Fisher Scientific). Endogenous biotinylation was assessed using a Streptavidin alone control.

### Statistical Analysis

Data sets were analyzed using Prism6 Graphpad. Where appropriate, Two-way ANOVA was utilized with multiple comparisons with no matching and Fisher’s LSD. Data are presented as ±SEM and differences are considered significant when *=p ≤ 0.05, **=p ≤ 0.01, ***=p ≤ 0.001, and ****=p ≤ 0.0001.

## Results

### Disruption of PaNA does not alter growth dynamics, but promotes pyocyanin production

We initially observed that PAO1Δ2794, a strain with an allelic deletion of PaNA, exhibits increased pyocyanin production compared to its wildtype counterpart, PAO1[44]. To determine whether this increased pyocyanin production was due to a disruption in PaNA, the enzyme which cleaves pseudaminic acid, we repeated the experiment with a PaNA transposon mutant, PW5679[33, 34]. Compared to its wildtype counterpart, MPAO1, again the PaNA mutant exhibits increased pyocyanin production (Figure 1A). As pyocyanin production is one QS readout, this observation suggests that PaNA and/or pseudaminic acid is important for QS. Both PaNA mutant strains exhibit altered behavior over the course of growth. That is, they both produce pigmentation, causing a shift in the yellowish media to a green that deepens over time. The strains begin producing this visible coloration at approximately six hours of growth. Pyocyanin was measured in the culture supernatants, as shown in Figure 1A. Both PaNA mutant strains produce significantly more pyocyanin under batch culture conditions than the wildtype strains. While the wildtype strains are capable of producing pyocyanin, under these growth conditions they produce nominal amounts that generally do not cause a visible color change in the culture media. Because QS is population density dependent, we next measured bacterial growth (Figure 1B). Figure 1B shows the growth of the PaNA mutant strains over 12 hours compared with wildtypes PAO1 and MPAO1. Population density was measured as the Optical Density (OD600) over time. The mutant strains and wildtype strains grow in almost identical patterns indicating that the loss of PaNA does not affect the bacteria’s viability or ability to grow in batch culture.

**Fig 1.**
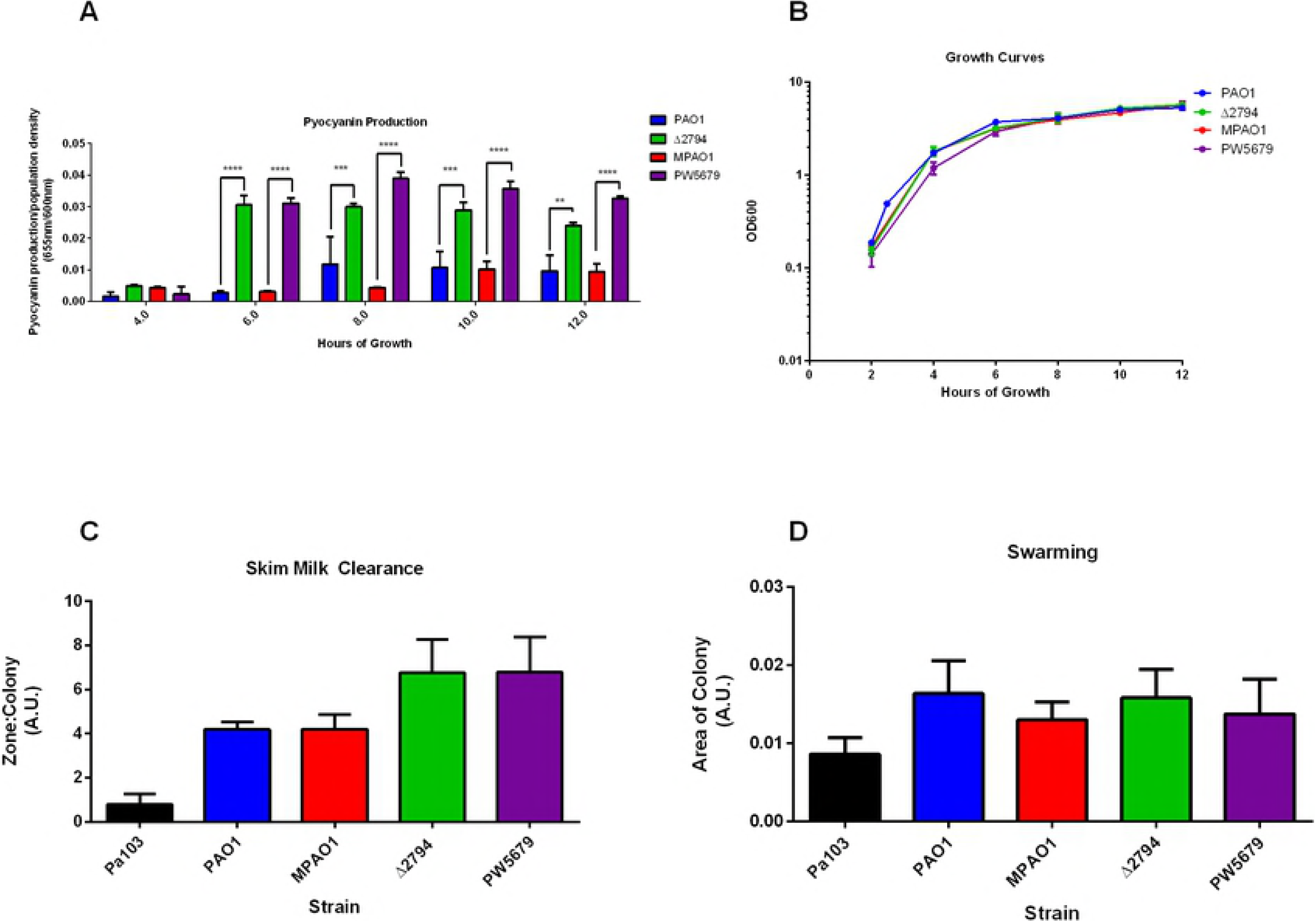
Disruption of PaNA elicits enhanced pyocyanin production. PaNA mutants produce more pyocyanin and sooner in the growth curve than wildtype populations. The represented data are an average of an n=3 where the mutant is compared to its corresponding wildtype at that timepoint. Significance is indicated when *p≤0.05, **p≤0.01, ***p≤0.001, and ****p≤0.0001. (1A). Wildtype strains and mutant strains exhibit comparable growth in LB media (n=3) (1B). There was a trend towards increased Las activity in the PaNA mutants (n=3) (1C), however, there was no difference from wildtype in swarming behavior (n=3) (1D), indicating that pyocyanin is the most affected level of QS.

### Disruption of PaNA does not alter LasB production or Swarming Behavior

Pyocyanin production is largely controlled by the PQS branch of QS, however, the PQS branch itself is regulated by the Las system[9]. We next asked whether the disruption of PaNA caused increased activity in all levels of QS. Skim-milk clearance was utilized to assess activation of the Las system as the Las-controlled LasB enzyme will degrade casein in the milk producing a clear zone around the colony. Nutrient agar plates containing skim milk were inoculated with the strains and allowed to grow overnight. After 24 hours of growth, the plates were photographed and ImageJ was utilized to measure the area of the zone of clearance, the area of the colony, and the area of the plate. The ratio of the zone of clearance to the colony size normalized by the area of the plate is shown in Figure 1C. While there is no significant difference between the PaNA mutants and their respective wildtype controls, there is a trend towards an increase in the area of clearance in the PaNA mutants. This experiment was controlled using a naturally LasB deficient wildtype strain, Pa103[48].

We next interrogated the Rhl branch of QS using swarming motility of the bacteria as an indicator of rhamnolipid production. *P. aeruginosa* can exhibit swarming motility on soft agar plates under appropriate conditions. This behavior relies on multiple factors including functioning flagella, but also the production of rhl-regulated rhamnolipid. Rhamnolipid acts as a biosurfactant to lower the surface tension of the agar allowing the bacteria to swarm away from the inoculation site. This experiment was controlled with a naturally non-flagellated strain, Pa103[49]. No mutant strain exhibited increased swarming motility compared with its parental wildtype strain. However, as all the strains, with the exception of Pa103, exhibited swarming, it also indicates the flagella of the PaNA mutant strains are functional and able to allow this form of motility (Figure 1D). Taken together, these data indicate that despite nearly identical growth patterns and population density, disruption of the PaNA gene causes an anomalous QS phenomenon, namely the over-production of pyocyanin.

### Pyocyanin production can be induced in the wildtype strains using conditioned media

We next asked whether this decoupling of pyocyanin production in the mutant strains from population density was in fact a QS-regulated phenomenon. In order to address this we conducted an induction experiment based on a central tenant of QS: the signal should be transferable and the behavior inducible at an earlier timepoint in a wildtype population[50]. Early work in QS showed that the quorum, or the minimal behavioral unit of bacteria, could be left-shifted using conditioned media which would contain the signaling molecules necessary to potentiate the behavior[5, 50, 51]. We grew wildtype PAO1 and MPAO1 in the presence of 10% culture supernatant harvested from overnight culture of the wildtype strains and the PaNA mutant strains. We then measured the pyocyanin production of the wildtype strains over 24 hours. Figure 2 shows that the signal is transferable to the wildtype strains and the behavior is inducible in PAO1 (Figure 2A) and MPAO1 (Figure 2B).

**Fig 2.**
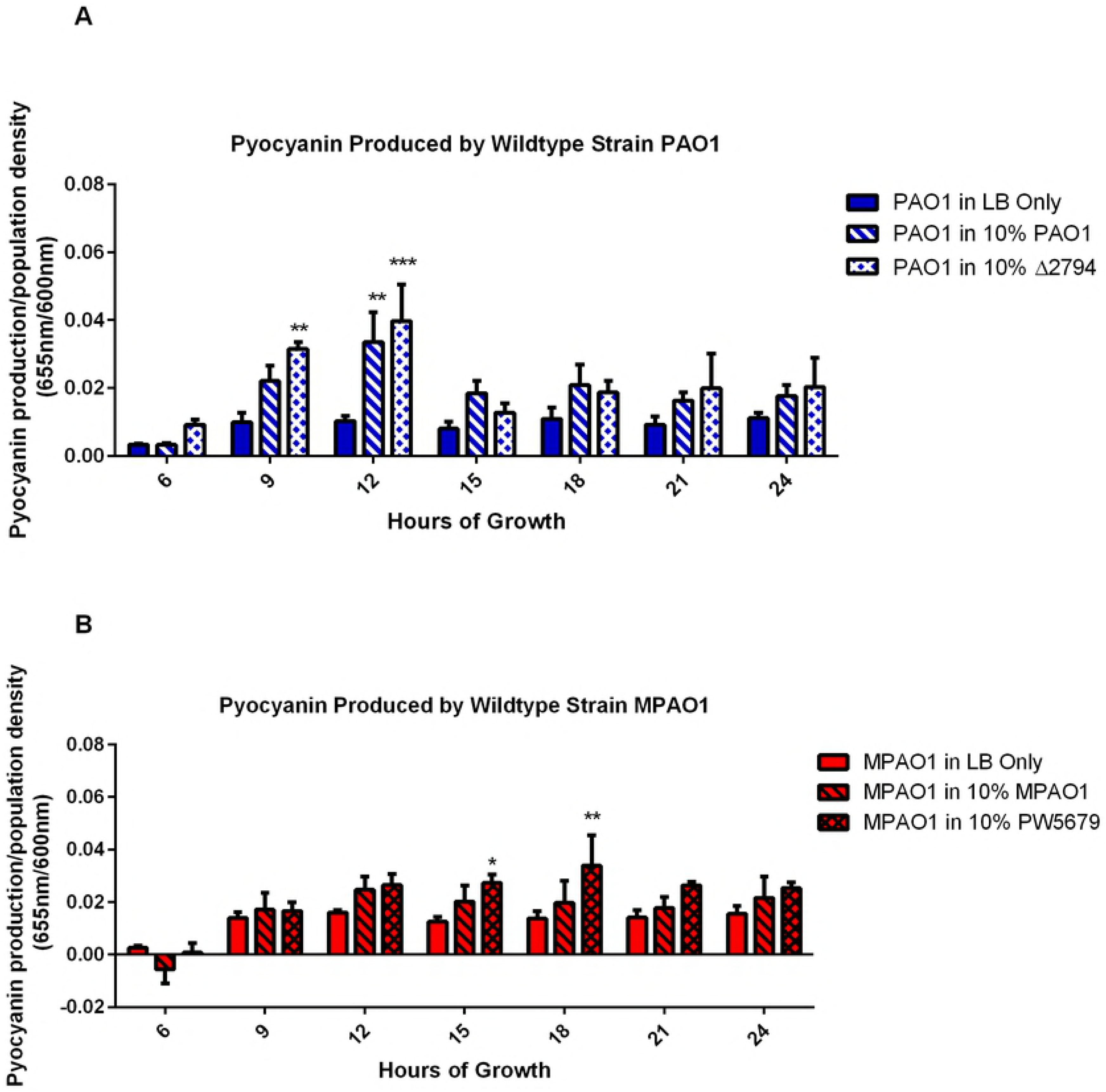
Wildtype strains can be induced to produce pyocyanin in conditioned media. Pyocyanin is produced in the wildtype strains in response to growth in the presence of 10% conditioned media indicating transferability of the QS signaling molecules. Wildtype strain PAO1 (2A) was grown with 10% cell-free culture supernatant from PAO1 and PAO1Δ2794 (n=3). MPAO1 (2B) was grown in 10% cell-free culture supernatant from MPAO1 and PW5679 (n=3). The represented data are an average of an n=3 where the mutant is compared to its corresponding wildtype at that timepoint. Significance is indicated when *p≤0.05, **p≤0.01, ***p≤0.001, and ****p≤0.0001.

Importantly, while the behavior is inducible earlier in wildtype strains grown in 10% supernatant, these cultures also produced pyocyanin at a higher magnitude than wildtypes grown in naïve media. The wildtype strains in naïve media do not exhibit an increased production of pyocyanin within the 24 hours assessed. The pyocyanin production in the wildtypes in naïve LB broth is relatively steady after 9 hours of growth and is rarely enough to cause a visible shift in the color of the media, while the wildtypes in media containing 10% culture supernatants produced increased pyocyanin production by 9 or 12 hours that is consistently higher than un-induced wildtypes. This increase in the pyocyanin production under the induced conditions, along with the occurrence at an earlier time point, suggests a left-shift in the quorum due to the artificial abundance of QS signal, indicating that this is a QS controlled phenomenon.

### Pyocyanin production relies on MvfR

We next asked whether pyocyanin production may be occurring through a pathway other than the canonical PQS system of QS. To address this question, we used an available mutant strain PW2812 which carries a disruption in the *MvfR* gene. As expected, culture supernatant from the PaNA mutants, wildtypes, or PW2812 was not able to induce pyocyanin production in the *mvfR*-mutant strain (Figure 3A). However, culture supernatant from PW2812 was able to induce QS in the wildtype MPAO1 (Figure 3B), indicating that the las and rhl signals are present in the PW2812 strain. Taken together, we determined pyocyanin production is reliant on functional MvfR. This further supports the indication that the disruption of PaNA impacts the PQS arm of QS to cause an overproduction of pyocyanin.

**Fig 3.**
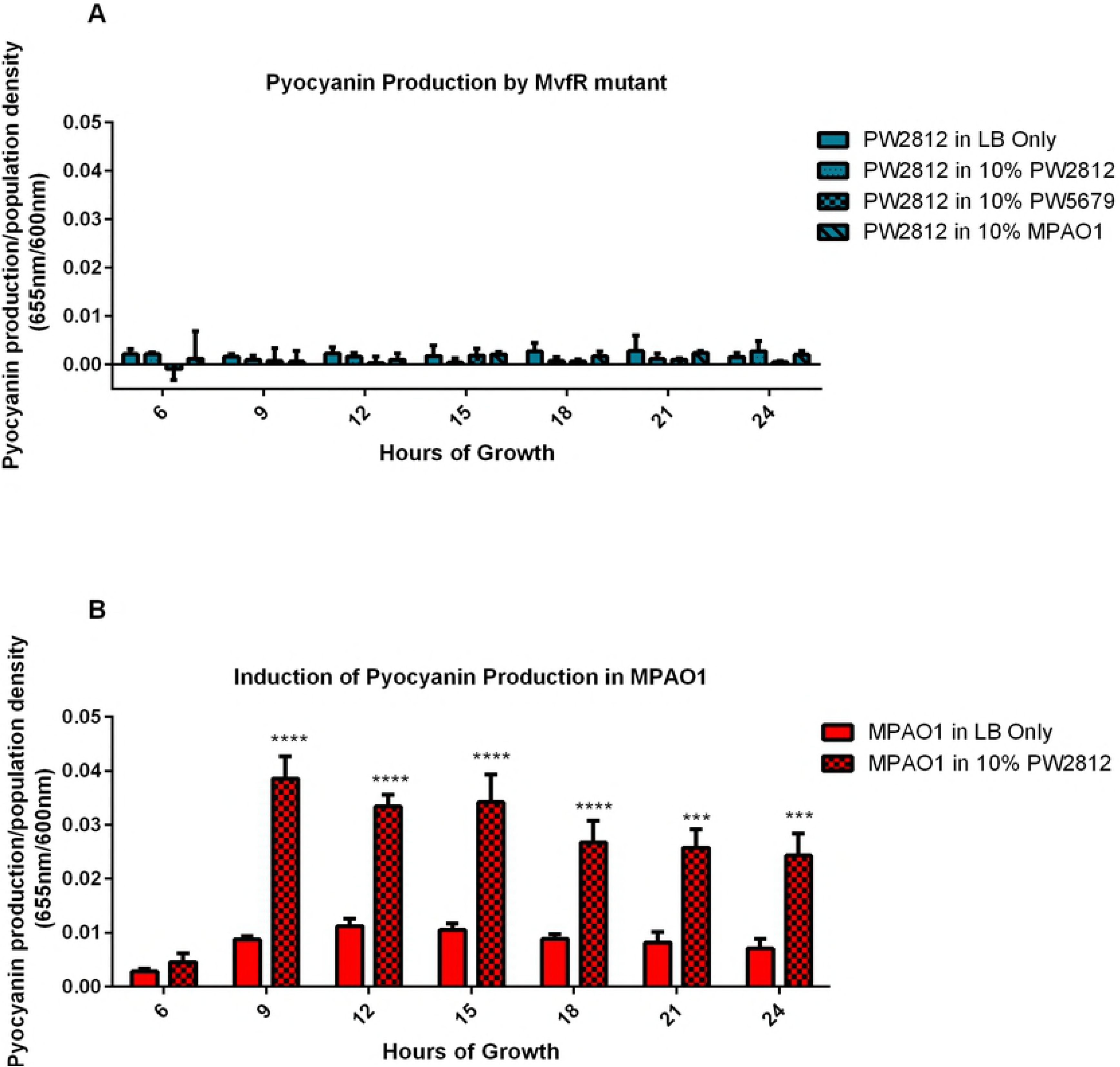
Pyocyanin production is MvfR dependent. Strain PW2812, which harbors a disrupted *MvfR* gene, was incapable of producing pyocyanin in any conditioned media (n=3) (3A). However, conditioned media from PW2812 was able to induce QS in the wildtype MPAO1 (n=3) (3B). The represented data are an average of an n=3 where the mutant is compared to its corresponding wildtype at that timepoint. Significance is indicated when *p≤0.05, **p≤0.01, ***p≤0.001, and ****p≤0.0001.

#### Glycosylation is temporally dynamic and alters during QS

We next asked three questions: 1) Are there differences in protein glycosylation patterns between the PaNA mutants and wildtype? 2) Do glycosylation patterns change in the wildtype strains over time? and 3) Do glycosylation patterns in the wildtype change during QS? In order to address these questions, we conducted lectin blots on lysate fractions from bacteria grown either in native LB or after induction to quorum sense. Figure 4 shows the dynamic nature of glycosylation patterns as evidenced by the binding of fucose-specific lectin from Lotus. This lectin was chosen because there were indicators that fucose may be a likely sugar to detect. Namely, the *P. aeruginosa* lectin LecB is specific for fucose and has been shown to associate with the bacterial surface after it is secreted [52]. Figure 4 shows the patterns of glycosylation change both over the growth of the bacteria (4A) and in response to the induction of QS to produce pyocyanin (4B). We highlighted the molecular weight region above 75 kDa due to minor endogenous biotinylation observed below 75 kDa in a streptavidin-only control blot. Despite limiting the view of the blots, this area shows that glycosylation is dynamic over time as well as is altered during QS in the wildtype. Specifically, a doublet develops around 100 kDa when MPAO1 is induced to QS (Figure 4B). Additionally, this doublet also appears by 24 hours of growth in the MPAO1 strain grown in naïve LB broth (Figure 4A). This indicates that glycosylation is a dynamic occurrence in the wildtype strain and is a responsive feature of the physiology of the bacteria. Additionally, we asked whether there were inherent differences in the patterns of glycosylation between the wildtype strains and the strains carrying mutations in PaNA. Using fucose-reactive Lotus lectin, Figure 4C shows that there are differences in protein glycosylation between wildtype strains and PaNA mutants. The 100 kDa doublet seen in QS-induced MPAO1 is present at 6 hours in both PaNA mutant strains but is absent from both MPAO1 and PAO1 at 6 hours. Additionally, there is a molecular weight shift in a pair of bands occurring at approximately 75 kDa, wherein the bands in the wildtype migrate slightly further down the gel than do the bands in the mutant strains. Taken together these data show that glycosylation, as represented by fucosylation, is dynamic during the growth of the bacteria, is altered in the PaNA mutants, and is altered in QS. It will be important to determine the nature of the glycan and the identity of the proteins involved. However, it is beyond the scope of this work.

**Fig 4.**
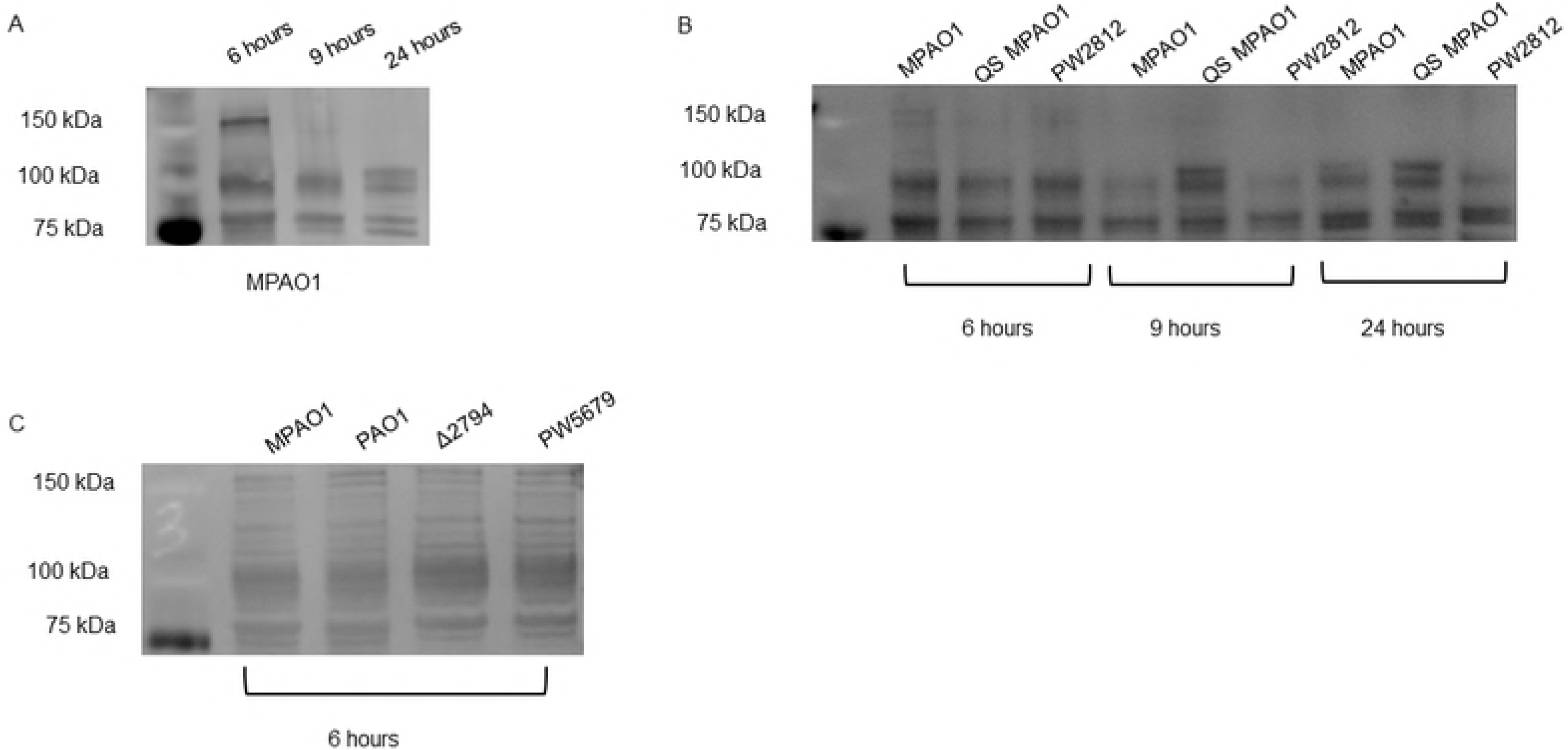
Glycosylation of *P. aeruginosa* is dynamic. Figure 4A shows that the lectin-binding pattern of fucose-specific Lotus lectin is dynamic over growth in wildtype MPAO1 (n=3). Figure 4B shows that there are differences in the glcosylation pattern of MPAO1 when it is induced to QS by growth in 10% culture supernatant from PW2812. PW2812 is included as a control, but also shows differences from wildtype. Namely a doublet at ~100 kDa does not develop in PW2812, even by 24 hours. The QS-induced MPAO1 develops a doublet by 9 hours whereas the naïve MPAO1 develops the double by 24 hours (n=3). Interestingly the doublet is present at 6 hours in the PaNA mutants (Figure 4C). In addition to the doublet, there is a molecular weight shift in a pair of bands occurring at 75 kDa. The bands in the PaNA mutant are slightly higher in molecular weight than the bands in the wildtype MPAO1 and PAO1 strains (n=3). Blots shown are representative of n=3 independent experiments.

### Disruption of multiple glycosyl-associated enzymes results in increased pyocyanin production

We next asked whether this observed dysregulation of pyocyanin production, putatively via a dysregulation of the QS arm PQS was limited to disruption of PaNA or whether it may be extrapolated to a global physiological relevance of glycosylation. We addressed this question by using an assortment of strains carrying transposon insertions of glycosyl-associated genes (Table I). Some of these disruptions were made in glycosyl-hydrolases, and others were made in glycosyl-transferases. Many of these are classified as “probable” proteins[32-34]. Consistent with the growth of the PaNA mutants, Figure 5A shows that all the strains grow approximately the same under batch culture conditions. Figure 5B and 5C show the pyocyanin production relative to population density of the glycosyl-transferases and glycosyl-hydrolases, respectively. In nearly all strains, there is an increased production of pyocyanin relative to the parent strain occurring at approximately 6 hours of growth. The only strain that failed to achieve significant pyocyanin production was PW3528, which contains a mutation in PA1391 encoding a probable glycosyl-transferase. This uncharacterized protein is proposed to be a glycosyl-transferase based on a conserved structural motif, however, it has had no demonstrated function[32]. While this strain, like many others used in this work, has an uncharacterized function, it’s lack of pyocyanin production suggesting a degree of variability in the roles that glycosylation plays in QS, but also importantly indicates that the phenomenon is not artifact of the mutation strategy. The observation of increased pyocyanin production in the majority of glycosyl-associated mutants is in agreement with what we observed in the PaNA-disrupted strains. Likewise, we again assessed Las activation with the skim-milk clearance assay (Figure 5D). Some strains exhibited increased clearance compared with the wildtype strain, while others did not. The assessment of rhl activation by use of the swarming assay showed no difference among any of the strains in the ability to swarm (Figure 5E). This ability to swarm is indicative of functioning flagella as this experiment was controlled with the Pa103 strain which does not possess flagella[49]. This is significant in that the flagella is a site known to be glycosylated in some strains of *P. aeruginosa* which may affect motility[53]; however, the flagella appear to be functional in all of the glycosyl-associated mutant strains.

**Fig 5.**
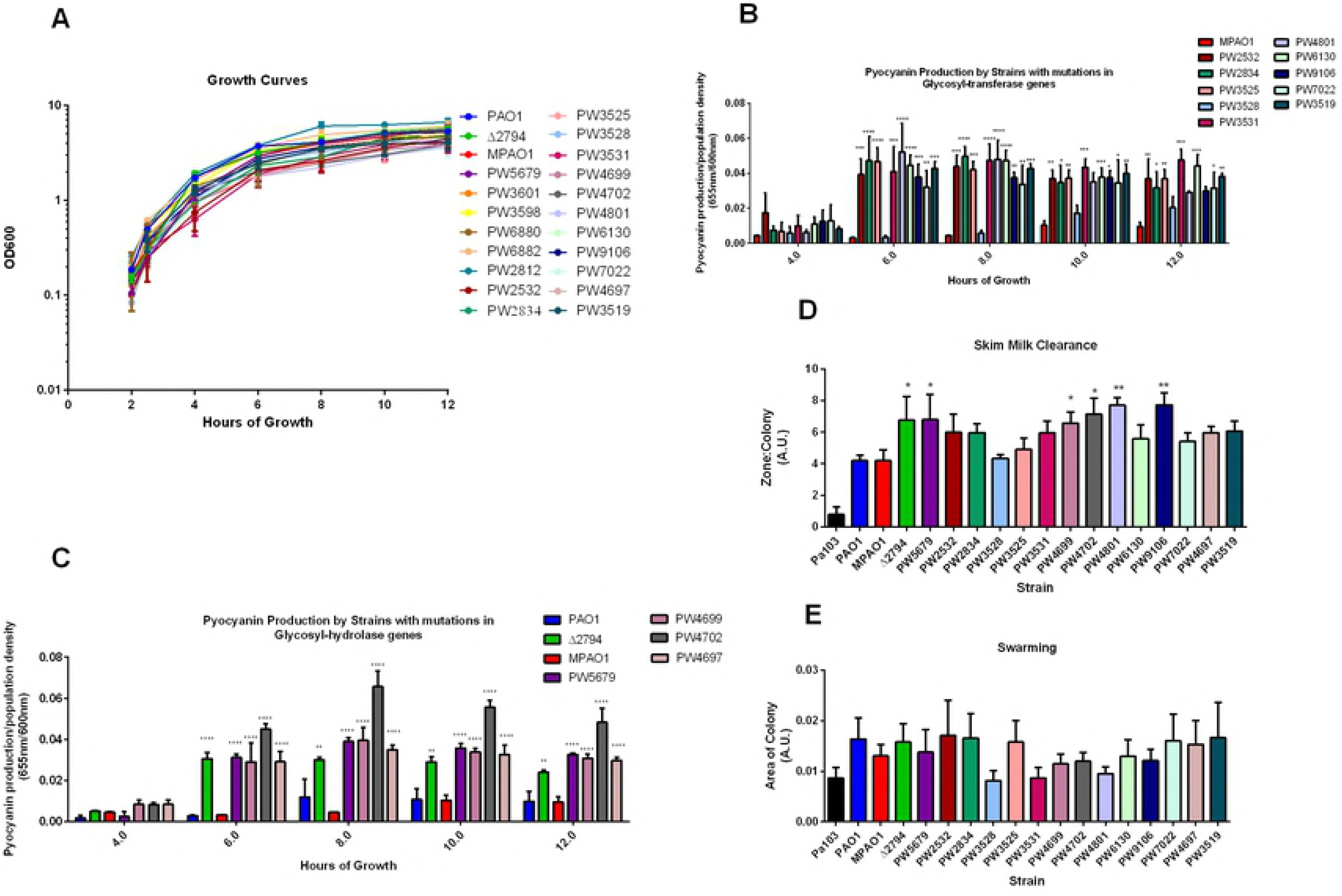
Enhanced pyocyanin production is a global consequence of disruption of glycosyl-associated enzymes. All strains grow equally well in LB broth (n=3) (5A). Nearly all strains carrying a disrupted glycosyl-transferase (5B) or glycosyl-hydrolase (5C) produced more pyocyanin, sooner than wildtype (n=3). There was a trend towards increased Las activity (5D), however, there was no difference from wildtype in swarming behavior (5E), indicating that pyocyanin is the most affected level of QS (n=3). The represented data are an average of an n=3 where the mutant is compared to its corresponding wildtype at that timepoint. Significance is indicated when *p≤0.05, **p≤0.01, ***p≤0.001, and ****p≤0.0001.

Finally, we assessed whether the signal was transferable from some of these glycosyl-associated mutant strains to wildtype strains. We included two strains with disrupted QS-regulator enzymes as a control[33, 34]. Culture supernatants were able to induce pyocyanin production above the background wildtypes in naïve culture media (Figure 6). Even at 24 hours, the pyocyanin produced by induction was detected at a higher magnitude than that of the uninduced wildtypes.

**Fig 6.**
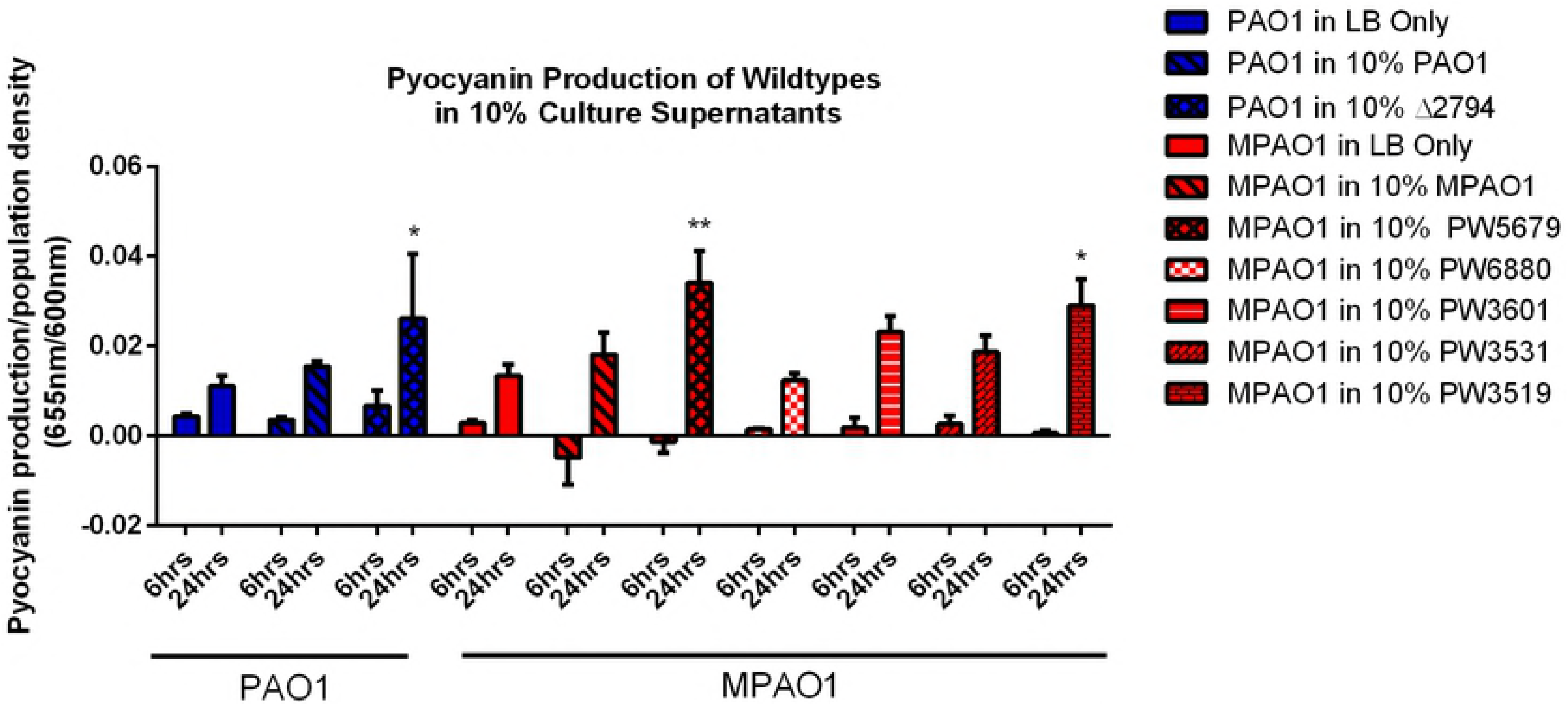
Wildtype strains can be induced to produce pyocyanin in conditioned media from various strains. Pyocyanin is produced in the wildtype strains in response to growth in the presence of 10% conditioned media indicating transferability of the QS signaling molecules. We assessed a few glycosyl-associated enzymes and included two strains containing transposons in QS regulators. Even at 24 hours there remains significantly increased levels of pyocyanin produced by wildtypes when induced to QS than produced in naïve media (n=3). The represented data are an average of an n=3 where the mutant is compared to its corresponding wildtype at that timepoint. Significance is indicated when *p≤0.05, **p≤0.01, ***p≤0.001, and ****p≤0.0001.

## Discussion

This research began with the observation that a strain of *P. aeruginosa* PAO1 from which the neuraminidase (PaNA) gene had been deleted, PAO1Δ2794, exhibited an altered phenotype from its parental wildtype. Namely, at equal population density, the deletion mutant produced visible pigmentation under normal growth conditions while the wildtype did not. This observation led us down the path of an investigation of the role of PaNA, its target pseudaminic acid/sialic acid, and overall glycosylation in QS of *Pseudomonas aeruginosa* PAO1. Growth curves conducted revealed that wildtypes and mutants grow approximately the same in LB broth at 37°C with shaking. This indicates that the disruption of PaNA as well as the other mutations explored do not impact the viability of the bacteria. That is, they are apparently equally capable of utilizing the nutrients available in LB broth to multiply and perform functions of life in a pure culture. Interestingly, the PaNA mutants along with most other glycosyl-associated mutants do produce visible pigmentation—typically around six hours of growth. This hyper-pigmentation is not observed in strains carrying mutations in the QS regulator genes with the exception of the LasI mutant. It was found also that culture supernatant from glycyosyl-associated mutant strains grown overnight could induce a hyper-pigmentation in the wildtype strains. This indicates that the signal molecules of the QS cascade are present in the supernatant and are able to initiate and potentiate the QS phenotype in the wildtype strains meaning that the signal is transferable. Transferability of the signal to induce the behavior implicates a QS phenomenon[50]. Notably, we observed a delay in the production of pyocyanin in the induction experiments. In the mutant strains grown in LB broth alone, visible pyocyanin is produced by six hours of growth, however in the induction assays, the earliest production of pyocyanin by wildtypes is around 9 hours of growth. This could be due to a less robust response of the wildtypes compared to the mutants. That is there are simply not enough bacteria making pyocyanin so it does not accumulate in the media as quickly. Alternatively, this could speak to a difference in the signal generated in the mutants and/or the response to that signal.

One observed phenomenon during the course of this study is that over time the magnitude of pyocyanin measured in the culture supernatants relative to the population density diminishes. Possible explanations of this include the complexity of the regulation of pyocyanin production and/or the natural cycling of pyocyanin. It is possible that at later growth phases the bacteria begin to turn off the production of pyocyanin, although pyocyanin production is generally associated with stationary growth[54, 55]. Pyocyanin production is largely under the control of the PQS system of QS, which itself is regulated by the Las system to express MvfR, as well as PqsH which is involved in the final conversion of the quinolone HHQ to PQS [6, 7]. Both HHQ and PQS are capable of binding to MvfR and potentiating their production, however, *in vitro,* PQS appears to bind more robustly [7]. Aside from the QS level regulation resulting in the production of the pyocyanin precursor PCA from two identical, but not entirely redundant, phenazine operons that are differentially regulated by HHQ-activated and PQS-activated MvfR, there are three PCA modifying enzymes that produce pyocyanin. [56, 57]. These enzymes are also under the complex regulation of a two-component nutrional regulator CbrA/CbrB [54]. Changes at any point in this pathway could alter the production of pyocyanin. Another explanation is that the pyocyanin produced is being reduced to another form not detected at the wavelength measured. Pyocyanin can be neutralized by two-electron reduction to a colorless product, or by glutathione to a less potent red-brown pigment[58, 59].

The present work shows that disruption of various glycosyl-associated enzymes leads to an over-production of pyocyanin. This apparent de-coupling of population density and QS in the PQS branch is a novel finding. Further, glycosylation has not previously been linked to QS. We conducted lectin blots on lysates of bacterial pellets that were collected either during the induction experiment using the *mvfR* mutant PW2812 or independently for the purpose of lectin blotting. Lectin blots with fucose-specific lectin from Lotus revealed three novel pieces of information. First there was a difference in the protein glycosylation pattern of the PaNA mutants compared to wildtype. This confirmed preliminary data that initiated this investigation. Secondly, the glycosylation pattern of the wildtype MPAO1 is dynamic over time and growth. Thirdly, the induction of MPAO1 to QS by growing the bacteria in the presence of 10% culture supernatant from PW2812 revealed that there are alterations in the glycosylation patterns that correspond with a QS phenotype. The induction experiment with PW2812 was used for lectin blotting because PW2812 contains a mutation in a QS regulator, not a glycosyl-associated enzyme. While we recognize the importance of identifying the proteins and glycans involved in this pathway, it is beyond the scope of this work. Overall, the observation of glycosylation changes in the induced wildtype samples strengthens the argument that wildtype glycosylation is an important determinant of QS.

We used a fucose-specific lectin from Lotus due to various indications that at least some proteins may be fucosylated at the cellular surface. Additionally, *P. aeruginosa* produces a fucose-specific lectin, LecB with has been shown to have functional roles in biofilm[52]. Fucose is an intriguing sugar that may be a component of either O-linked or N-linked glycans[60]. Fucosylation has been demonstrated as an important determinant of the microbiome and colonization of the gut, but this is largely in relation to fucosylation occurring on gut mucins[61]. However, fucose can be utilized as a carbon source for metabolism and can affect behavior of bacteria[62]. Our interest, however, lies in the occurrence of fucose as a structural modification and/or signaling mechanism. Altering the composition of the glycan may change the conformation of the glycan and or protein[63]. Post-translational modifications that alter the conformation of the protein can have profound consequences to signaling cascades. This is well understood for phosphorylation of proteins, however, has not been well explored in glycobiology of *P. aeruginosa.* We postulate that glycosylation may act as signaling mechanism that regulates QS.

Our knowledge of protein glycosylation in *P. aeruginosa* is limited to a few characterized proteins and glycans: namely the glycosylation of pili, flagellin, and LecB lectin[14, 17, 18, 23-28, 53]. One possible target of interest is the outer membrane porin, OprF. It has been reported to interact with the lectin, LecB, which recognizes fucose residues[14, 52]. Additionally, OprF has been reported to stimulate QS behavior in the bacteria in response to activation during infection by host INFƔ[64]. OprF provides an interesting juxtaposition of being an outer membrane protein that responds to the environment, likely glycosylated as evidenced by its interaction with LecB, as well as a demonstrated role in QS regulation[52, 64-66]. While this work does not address any specific glycan or protein, it does highlight a physiological phenomenon that is largely unexplored. As evidenced by this work, this has profound consequences for the bacteria. Considering the importance of pyocyanin as a virulence factor, understanding the sequence of events that leads to its over-production is vastly important. Beyond pyocyanin alone, the novel finding that QS may be dysregulated when glycosylation is dysregulated has a plethora of opportunities for future studies. Quorum sensing plays important roles in the bacterial physiology and life cycle and is an important component of *P. aeruginosa’s* ability to cause infection or establish a mature biofilm[67-75]. Understanding the roles of glycosylation in QS in *P. aeruginosa* may very well open new avenues of research for developing strategies for combatting infections caused by *P. aeruginosa*. This work lays the groundwork for exploring glycosylation as an important regulatory system which could be a therapeutic target.

This work also advances the field of glycobiology in *P. aeruginosa* by presenting a preliminary exploration of uncharacterized probable glycosyl-associated enzymes. A few of the strains used carry mutations in enzymes that have been described in various glycosylation-associated functions such as exopolysaccharide construction[32-34, 44]. However, the majority of the enzymes explored have unknown functions and unknown substrates: both carbohydrate and platform such as protein or lipid[32-34]. We show, however, that these enzymes have some physiological significance to the bacteria. We provide a foundation for further exploration of these enzymes to gain understanding of how they interact with QS as well as to characterize their function and substrates. Additionally, this work shows that even the enzymes which have a characterized function may play multiple roles in the bacteria. For example, strains PW6130 and PW4801, which have mutations in genes associated with biofilm exopolysaccharide synthesis, also exhibit an over-production of pyocyanin[32-34]. While we did not explore the biofilm formation in these strains, it is of note that disruption of PA2794 was described previously to inhibit the maturation of biofilm by PAO1[44]. It was also demonstrated that the deletion of PA2794 altered the virulence of the bacteria in a mouse model of infection[44]. Altered virulence and biofilm formation in these strains would indicate that these glycosyl-associated enzymes, and therefore glycosylation, is important for the viability and fitness of the bacteria. This raises the potential of the pathways and enzymes being therapeutic targets for combatting colonization and infection by *P. aeruginosa.*

Taken together this work has demonstrated that disruption of various enzymes associated with or predicted to be associated with glycosylation pathways of *Pseudomonas aeruginosa* leads to an over-production of pyocyanin, suggesting that proper glycosylation of bacterial proteins is critical for QS regulation. This work underscores the importance of broadening our understanding of the role of glycosylation in QS regulation, and we have provided a foundation for a rational exploration of the field.

## Acknowledgements

We would like to acknowledge Viktoriya Soludushko and Dr. Barnita Haldar for their advice and technical assistance, Dr. Alice Prince for the generous gift of PAO1 and PAO1Δ2794 strains, Dr. Jonathan Audia for advice and the gift of Pa103 strain as well as Drs. Troy Stevens, Diego Alvarez, and Paul Brett for their guidance on this research.

